# Trait-trait relationships and tradeoffs vary with genome size in prokaryotes

**DOI:** 10.1101/2021.07.23.453341

**Authors:** Sara Beier, Johannes Werner, Thierry Bouvier, Nicolas Mouquet, Cyrille Violle

**Affiliations:** Leibniz Institute for Baltic Sea Research Warnemünde (IOW), Department of Biological Oceanography, Rostock, Germany; UMR 7621 Laboratoire d’Océanographie Microbienne, Observatoire Océanologique de Banyuls-sur-Mer, Sorbonne Université, Banyuls-sur-Mer, France; High Performance and Cloud Computing Group, Zentrum für Datenverarbeitung (ZDV), Eberhard Karls University of Tübingen, 72074 Tübingen, Germany; MARBEC, Université de Montpellier, CNRS, Ifremer, IRD, Montpellier, France; FRB – CESAB, Montpellier, France; CEFE, Univ Montpellier, CNRS, EPHE, IRD, Montpellier, France

## Abstract

We report genomic traits that have been associated with the life history of prokaryotes and highlight conflicting findings concerning earlier observed trait correlations and tradeoffs. In order to address possible explanations for these contradictions we examined trait-trait variations of 11 genomic traits from ~ 18,000 sequenced genomes. The studied trait-trait variations suggested: (i) the predominance of two resistance and resilience-related orthogonal axes and (ii) at least in free living species with large effective population sizes whose evolution is little affected by genetic drift an overlap between a resilience axis and an axis of resource usage efficiency. These findings imply that resistance associated traits of prokaryotes are globally decoupled from resilience related traits and in the case of free-living communities also from resource use efficiencies associated traits. However, further inspection of pairwise scatterplots showed that resistance and resilience traits tended to be positively related for genomes up to roughly five million base pairs and negatively for larger genomes. This in turn may preclude a globally consistent assignment of prokaryote genomic traits to the competitor - stress-tolerator - ruderal (CSR) schema that sorts species depending on their location along disturbance and productivity gradients into three ecological strategies and may serve as an explanation for conflicting findings from earlier studies. All reviewed genomic traits featured significant phylogenetic signals and we propose that our trait table can be applied to extrapolate genomic traits from taxonomic marker genes. This will enable to empirically evaluate the assembly of these genomic traits in prokaryotic communities from different habitats and under different productivity and disturbance scenarios as predicted via the resistance-resilience framework formulated here.

## 1. Introduction

Prokaryotes contribute largely to the global organic carbon budget (Whitman et al., 1998), are main drivers of major element cycling (Konopka et al., 2015), and are therefore key compounds of earth functioning. However, natural microbial communities are typically extremely diverse and complex, and it remains challenging to predict prokaryote ecosystem functioning and community dynamics in response to environmental changes (Shade et al., 2012; Bardgett and Caruso, 2020). A large body of research highlights the impact of structural community properties such as diversity and species interaction patterns on community functioning (Poisot et al., 2013; Duffy et al., 2017). The effect of structural community properties, however, depends on the characteristics of individual species in a community mediated via their traits and the distributions of these traits have been shown to influence community functioning and dynamics (Enquist *et al*., 2015). Microbes harbor an enormous functional versality regarding the number of metabolic functions and pathways they are capable of, which poses a challenge in selecting meaningful functional descriptors to infer overall community functioning. To simplify the assignment of functional attributes to microbial communities it has been suggested to characterize communities based on the distribution of overall life histories rather than focusing on the potentially large number of traits related to specific metabolic pathways (e.g. Bardgett and Caruso, 2020; Malik *et al*., 2020). Life history traits determine how species allocate available energy among survival, growth and reproduction and are therefore decisive for the overall production and stability of an ecosystem.

Several traditional life history classifications allocate species into binary categories either concerning their response to environmental change or in dependence of the resource availability required for growth (Box 1). For instance, the characterization of species along the specialist-generalist continuum is related to their tolerance against environmental change (Kassen, 2002). It has been pointed out that traits that encompass the capability of species to tolerate or to adapt to changing conditions is related to *resistance*, i.e. the ability of organisms to withstand disturbances (Nimmo et al., 2015).

### BOX 1: Glossary

#### Species

The species concept developed for macroorganisms cannot be directly transferred to asexually proliferating microorganisms. However, sequenced genomes sharing >94% of their average nucleotide identity (Konstantinidis and Tiedje, 2005) or operational taxonomic units (OTUs) delineated e.g. from amplicon sequence variants (ASVs) of taxonomic marker genes approximate natural units that reflect ecological species and will be referred to as species.

#### Generalist/specialist

The characterization of species along the generalist specialist continuum is based on the organisms’ niche breadth, where ecological specialization indicates a limited niche breadth and niche breadth has been defined as the variety of resources, habitats, or environments used by a given species (Sexton et al., 2017). It has recently been pointed out that the unambiguous characterization of species along the specialist generalist continuum is challenging as niche breadth estimations usually refer to a specific range of measured conditions, and for instance, a resource generalists may simultaneously be a temperature specialist (Bell and Bell, 2021). Still, even though not practically measurable, the increasing number of biotic and abiotic conditions in a multivariate niche space under which a species can proliferate can be considered as an increasingly large multidimensional niche breadth of this species.

#### Colonizer/competitor

Competition-colonization tradeoff models assume that species can occupy a niche as colonizer by efficiently colonizing empty habitat patches or a niche as competitor by outcompeting species within sites (Tilman, 1994; Mouquet et al., 2005).

#### r/K selection

The theory of r- and K-selection postulates that r-strategists allocate an increased fraction of resources to reproduction under conditions of high density independent mortality (Gadgil and Solbrig, 1972).

#### Oligotrophic/copiotrophic

Oligotrophic species are able to grow in nutrient poor environments while copiotrophic species require high concentrations of inorganic and organic compounds (Poindexter, 1981).

#### CSR terminology

The CSR schema sorts species into competitors (C), stress-tolerators (S) and ruderals (R). Ruderals have an advantage in habitats with high disturbance levels, stress tolerators dominate habitats with high stress levels such as low nutrient conditions or extreme temperatures, while competitors profit from their competitive advantage over other organisms in resource-rich habitats (Grime, 1977).

#### Genomic trait

The term trait has previously been defined as ‘any morphological, physiological or phenological feature measurable at the individual level, from the cell to the whole-organism level, without reference to the environment or any other level of organization’ (Violle et al., 2007). Here, in order to adapt the use of “traits” to genomic data, genomic traits will be defined as variables that characterize prokaryotic species and which can be delineated from their genome sequence data, such as its genome size. We particularly focus on genomic traits that had been associated in earlier studies with one of the above listed life history traits.

The r/K selection theory refers to the fraction of resources allocated to reproduction (Gadgil and Solbrig, 1972). R-strategists can be described as opportunists that proliferate fast in response to opportune environmental changes while K-strategists are typically strong competitors that allocate more resources in efficient resource usage rather than growth. The competitor/colonizer classification (Tilman, 1994) differentiates between organisms with competitive advantage in either empty or densely populated habitats: as a consequence the best competitor will outcompete all other species at low disturbance rates, while species with high colonization efficacy will dominate ecosystems at high disturbance rates (Hastings, 1980). Both, r-strategists as well as species with high colonization efficacy should be characterized by short lag-phases and fast growth rates enabling them to respond fast to changing environmental conditions or rapidly colonize empty habitats. It has been proposed that traits that indicate rapid reproduction or strong re-colonization capabilities are linked to an organism’s *resilience*, defined as its capacity to recover after a disturbance (Nimmo et al., 2015).

Resistance and resilience related traits represent two distinct components of ecological units (i.e. populations, communities or ecosystems) that determine their response to disturbances (Shade et al., 2012; Nimmo et al., 2015). High levels of resistance and resilience are both associated with costs: Dall and Cuthill (1997) suggested costs that are inherent with a generalist’s life-style can include performance reductions from having more ecological variables to monitor. High potential growth rates in microbes often come at the expense of low resource usage efficiency (Fierer et al., 2007; Roller et al., 2016). In agreement with this, tradeoffs between resilience and resistance have been observed in microbes, where species can allocate transcriptional resources in the expression of stress resistance genes at the cost of a reduced expression of genes that promote growth (Ferenci, 2016). Such tradeoffs between resistance and resilience have been described not only at the species but also at the community level (e.g. de Vries and Shade, 2013; Garcia *et al*., 2020; Piton *et al*., 2021).

Resource availability can also shape the distribution of resistance or resilience related life histories due to the above mentioned potential costs that are linked to the response to disturbance. Traditionally, in microbial ecology, large emphasis has been put in sorting species according to the nutrient levels required for their growth, in either oligotrophic or copiotrophic species (Koch, 2001; Fierer, 2017). It has been suggested that copiotrophic species that require high nutrient levels for growth can commonly be characterized as generalists (Christie-Oleza et al., 2012) and / or r-strategists that react fast to nutrient pulses (Fierer, 2017). However, although some binary life history classifications may overlap, these terms should not be used interchangeably as they are defined differently (Box 1).

The more complex CSR theoretical schema, that goes beyond a binary classification of life-history traits, sorts species into three classes of ecological strategies (competitor vs. stress-tolerator vs. ruderal species) depending on their location along two major environmental gradients: disturbance and productivity (Grime, 1977). This framework was originally developed in plant ecology and integrates different dimensions in the interplay of resistance, resilience and resource availability. Several studies have suggested applying the CSR framework in the field of microbial ecology (e.g. Krause *et al*., 2014; Fierer, 2017). However, it has recently been claimed that the reliance of heterotroph microbial organisms on external carbon and energy sources distinguishes them from autotroph plants and complicates the application of the CSR framework to microbial communities dominated by heterotrophs. The CSR framework has therefore been adapted to microbial ecology by classifying high yield (Y) – resource acquisition (A) – stress tolerator (S) categories (Malik et al., 2020). Here Y refers to species with high carbon use efficiency, A replaces the plant competitor strategy because microbial competition is mainly over resources and the S strategy refers to species that are adapted to stress exposure due to deviations from ambient.

Although a number of prokaryote representatives from naturally abundant and relevant lineages have recently brought into culture (e.g. Marshall and Morris, 2013; Neuenschwander *et al*., 2018) the majority of prokaryotic species remains uncultured (Steen et al., 2019). Accordingly, the physiologically measured species characteristics that inform about the life-histories of prokaryotes are only available for a small minority of cultured representatives. However, a growing number of genome sequences from so far uncultured prokaryotic strains are available. In prokaryotes the myriad of responses to environmental change, such as changes in resource availability, abiotic stressors or biological interactions, is coded in their genomic material. Thus, the DNA of a population contains the information of its performance under all possible conditions or its coverage of the n-dimensional niche space. However, an incomplete understanding of the complexity of gene regulation from genome sequence data currently limits our capacity to predict gene expression patterns and consequently the specific phenotype of a strain in a given environment.

Still, it has been demonstrated via machine learning approaches that the presence of functional genes within genomes was tightly linked to the ecological niche occupied by the corresponding prokaryotes, explaining ~50% of their niche variability in a high-dimensional niche space (Alneberg et al., 2020). A more simplified possibility to characterize prokaryotes via their genomic information is to extract and evaluate simple parameters, so-called genomic traits, from their genomes, such as the presence or copy number of a specific gene or genome size. Although this option may lack accuracy due to the high degree of simplification, various studies found empirical and statistically validated evidence for correlations between certain genomic traits and the binary life history strategies listed in BOX1.

Hereafter, we summarize literature about genomic traits that were associated with statistical support to life history strategies with a focus on binary classifications and earlier reported correlation among these traits. In order to resolve some conflicting findings concerning earlier proposed genomic trait assignments to life histories, their pairwise correlations and tradeoffs, we inspected the covariation patterns among 11 genomic traits that are considered as life history proxies, from ~ 18,000 sequenced genomes representing ~9,000 species in a multivariable trait space. As a result, we categorized the studied genomic traits within a resistance-resilience based framework. This framework resolved conflicts and may be used in future studies to address community dynamics under different disturbance and productivity scenarios in variable habitats.

We demonstrated that all analyzed genomic traits featured significant phylogenetic signals and proposed a simple tool to empirically evaluate the theoretical predictions made here via the extrapolation of genomic traits based on taxonomic marker genes from uncultured taxa.

## 2. Genomic traits as life history proxies in prokaryotes

A number of earlier studies has addressed prokaryotic genome features that give indirect evidence for physiological characteristics and the life-history strategies of the corresponding organisms (Figure 1). Particularly the classification as generalist, measured as an organism’s physiological versatility or its ability to colonize multiple habitat types, has been related to a range of different genomic characteristics. This includes features such as a large genome size (Barberan et al., 2014; Bentkowski et al., 2015; Cobo-Simon and Tamames, 2017; Sriswasdi et al., 2017) and a high fraction of regulatory genes (Parter et al., 2007; Kostadinov et al., 2011). It indeed seems reasonable that organisms that are able to live and proliferate under variable conditions need more genes for sensing or for coping with a range of different growth conditions. They further need a larger number of transcription factors (%TF, i.e. regulatory genes) to regulate genes that are alternatively expressed depending on the prevailing environmental conditions. Based on the same considerations it has been suggested that a high frequency of genes acquired via horizontal gene transfer (%HGT) would increase the versatility of prokaryotes and allow them to grow in a larger number of environments (Takemoto, 2013).

**Figure 1:**
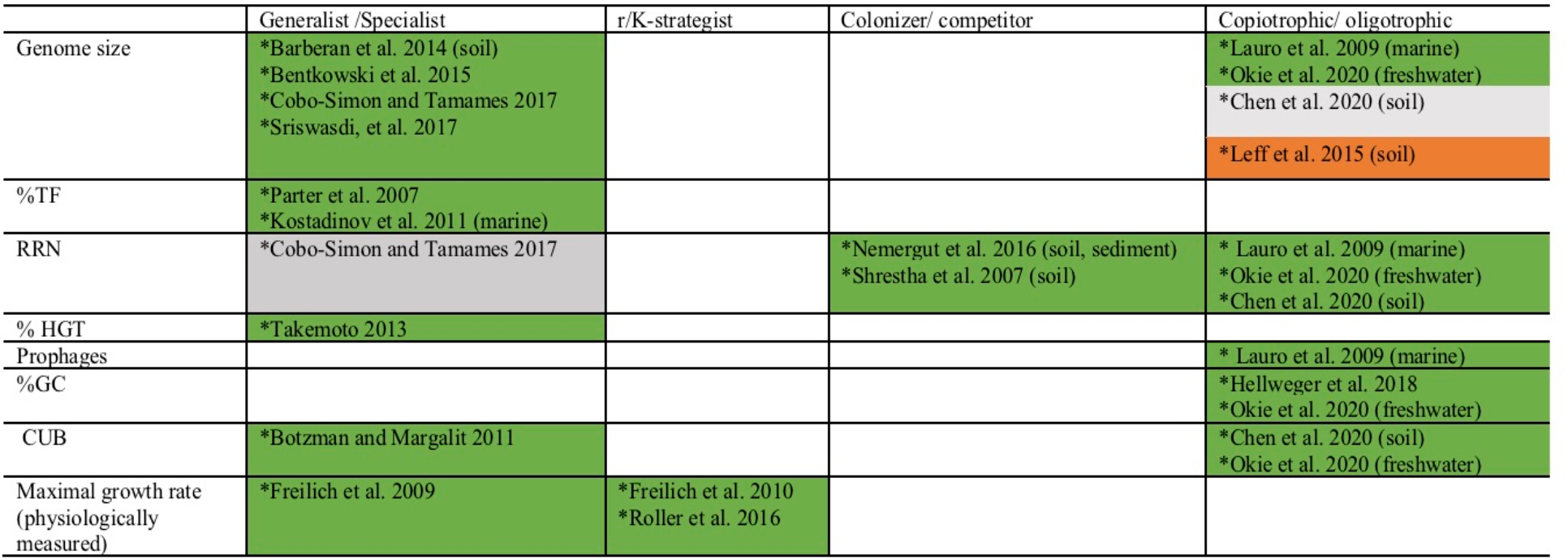
Overview of earlier literature that linked genomic traits or the physiologically measured maximal growth rates of prokaryotes to their life-history traits. If studies refer to organisms from a specific ecosystem, this is given in brackets. Green (or red) fields indicate studies that found positive (or negative) relationships (p < 0.05) between the listed genomic feature and a life style as generalist, r-strategist, colonizer or copiotroph. In one study no p-value was reported, but mathematical evidence for increased environmental tolerance of species with large genomes was provided via the computational simulation of evolutionary processes (Bentkowski et al., 2015). Gray fields indicates studies where neither a significant relationship and not even a positive or negative trend was detected (p>0.5).

It has further been argued that the life style as a generalist requires enhanced growth rates. In agreement with this, high growth rates (Freilich et al., 2009) as well as elevated codon usage biases (CUB, Botzman and Margalit, 2011) have been associated with a generalist life style. The CUB describes the phenomenon that synonymous codons are used unevenly among genes, where genes coding for highly expressed proteins are enriched in codons that reflect the taxon-specific tRNA pool. The CUB has been shown to be specifically pronounced in fast-growing organisms and was therefore interpreted as a genomic feature that correlates with maximal growth rates of prokaryotes (Vieira-Silva and Rocha, 2010).

In contrast, the number of rRNA gene copies (RRN) in prokaryote genomes could not be associated significantly to the classification as either generalist or specialist (Cobo-Simon and Tamames, 2017). Instead, the RRN decreased significantly during the community succession after environmental disturbances (Shrestha et al., 2007; Nemergut et al., 2016). A high RRN at early successional stages can be interpreted as evidence for a life style as colonizer and corroborates observations that associated high RRN with short lag phases and high growth rates (Klappenbach et al., 2000; Stevenson and Schmidt, 2004; Vieira-Silva and Rocha, 2010; Roller et al., 2016).

A significant correlation between growth rate and competitive ability (Klappenbach et al., 2000; Stevenson and Schmidt, 2004; Vieira-Silva and Rocha, 2010) or carbon use efficiency (Roller et al., 2016) demonstrated that prokaryotes can generally be well classified along the r/K-strategist continuum. Organisms with short duplication times and accordingly high CUB are therefore likely to be r-strategists.

It had been hypothesized that the evolution towards GC depleted genomes is an adaption to nutrient poor conditions because GC pairs contain one more nitrogen atom compared to AT pairs (Giovannoni et al., 2005; Grzymski and Dussaq, 2012). Hellweger and colleagues (2018) demonstrated that beside other mechanisms, such as mutation biases, particularly N limitation but also C-limitation impacted the evolution of genomes towards GC depletion. Accordingly, low GC content can be interpreted as a genomic trait that is indicative of an oligotrophic life style in prokaryotes. Alternatively, may a high frequency of GC pairs that have three hydrogen bounds compared to only two hydrogen bounds in AT pairs enhance the resistance of cells with high GC content against some specific stressors, such as heat or desiccation stress. Probably for this reason, a higher GC content was found in arid, nutrient-poor soils exposed to heat and dehydration stress than in soils with higher nutrient contents but lower stress exposure (Chen et al., 2020).

Lauro et al. (2009) suggested that high RRN, an elevated number of prophages and a large genome size are more common in marine copiotrophic than in marine oligotrophic bacteria. A recent study demonstrated in agreement with this that nutrient additions to oligotroph lake water induced a significant increase of community mean genome size, RRN, CUB and GC content (Okie et al., 2020). In contrast, nutrient additions to soil environments resulted in the selection of prokaryotes with smaller genomes (Leff et al., 2015) and a recent comparison of microbial communities in nutrient rich versus nutrient depleted soils did not reveal a significant difference in their average genome sizes (Chen et al., 2020). In addition, enhanced CUBs were found associated with copiotrophic microbial communities in soil environments (Chen et al., 2020).

## 3. Trait-trait correlations in the light of physiological constraints

A number of earlier studies explored pairwise correlations among the above-mentioned genomic traits or with the organism’s physiologically measured maximal growth rate. Most of these studies found significant positive correlations e.g. between genome size and %TF, RRN and growth rates (Figure 2). As a consequence, if large genome sizes and a high %TF are indicative of a generalist life history and a high RRN and elevated growth rates indicate r-strategists and/or colonizers, a generalist’s life style should be associated with a simultaneous life style as r-strategist and colonizer. This inference however contradicts the often observed and above outlined tradeoff between resistance and resilience in microbial communities (e.g. Ferenci, 2016; Garcia et al., 2020; Piton et al., 2021). A resistance-resilience tradeoff would instead imply a negative correlation between resistance associated traits, such as genome size or transcription factors versus resilience associated traits, such as rRNA gene copy numbers or growth rates. We want to point out that some studies did not find significant positive relationships between genome size and growth rates and although the observed correlation coefficients were positive, they were very weak (Vieira-Silva and Rocha, 2010; Westoby et al., 2021b). Still, also in these cases no negative correlations were detected as one would expect from a tradeoff between resistance and resilience.

**Figure 2:**
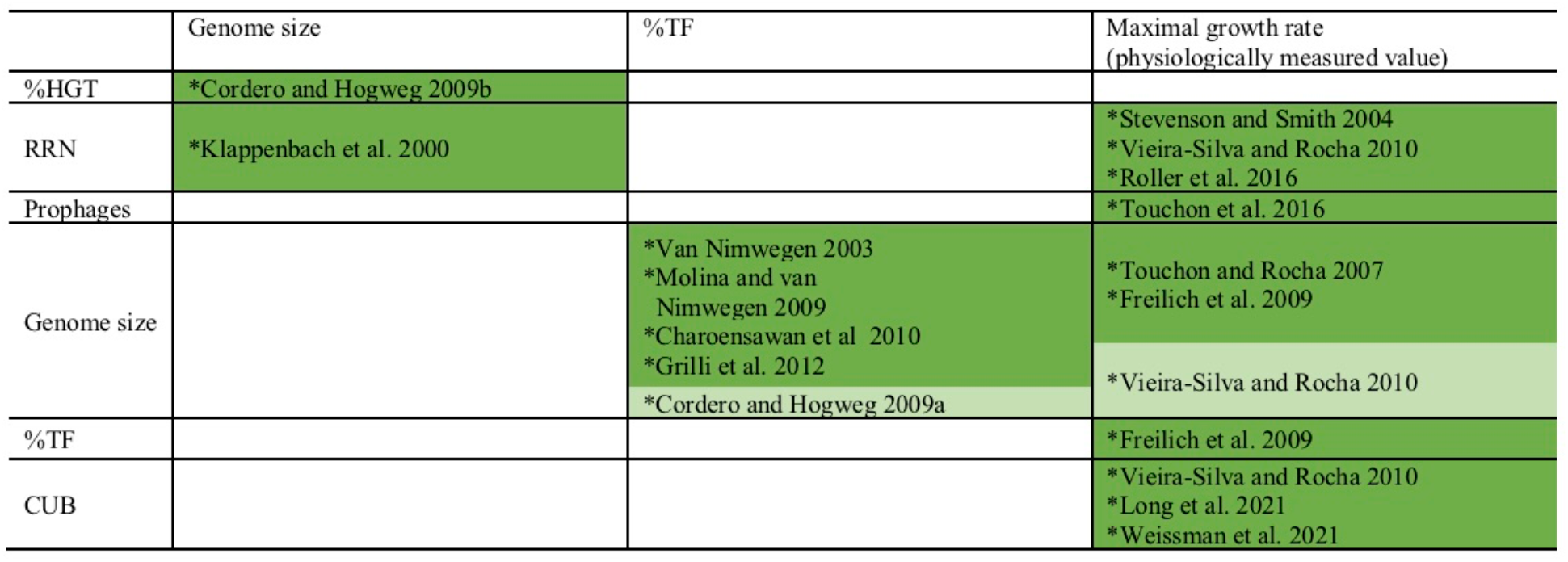
Earlier reported pairwise correlations among genomic traits and /or physiologically measured maximum growth rates. Dark green fields indicate positive significant correlations (p < 0.05) and light green field indicates a positive trend below the significance level (0.5 > p > 0.05).

In order to elucidating conflicting relationships between these earlier observations and the resistance-resilience tradeoff, we here examined covariations among multiple genomic traits. These traits were extracted from the genomic material of sequenced prokaryotic genomes available via the JGI/IMG platform (https://img.jgi.doe.gov/, Chen et al., 2021; Mukherjee et al., 2021). For downstream analyses we considered those genomes that are integrated into the default reference database in the PICRUSt2 software (Douglas et al., 2020) and which infers the genomic content of uncultured prokaryotes from closely related genomes via taxonomic marker genes. In total 17,856 of the 20,000 genomes that are integrated in the PICRUSt2 default reference phylogenetic tree could be downloaded via the JGI/IMG database (date of download: 24.08.2021). These genomes cover a broad phylogeny (73 phyla; 172 classes; 382 orders; 762 families; 2,669 genera; 8,847 species) and include genomes from yet uncultured candidate phyla (Table S1).

Additionally to the genomic traits presented above (Figure 1), we also considered genome level parameters for gene richness (i.e. the number of different genes in a genome) and the gene duplication level (i.e. the average number of gene copies per gene in a genome). This is because the genome size can increase due to the integration of new genes or due to the duplication of already existing genes and either one of these two mechanisms may have different consequences for the resistance level of prokaryotes: on the one hand may different copies of the same gene be expressed alternatively in response to environmental change, for instance when they differ in their pH optimum (Miyashita et al., 1991) and therefore contribute to resistance levels analogously to two different genes that are can be expressed alternatively. On the other hand may multi copy genes have an adaptive effect to stressful environment due to an enhanced dosage effect (Kondrashov, 2012). Some genomic traits are provided directly by the JGI genome statistics, while others were computed from the genome sequences. An inspection of the JGI/IMG provided RRN data indicated inaccuracies for this specific genomic trait (Supplementary Figures S1/S2). We therefore extrapolated RRNs for the JGI/IMG reference genomes affiliating with the same genus of genomes stored in the Ribosomal RNA Operon Copy Number Database (rrnDB, https://rrndb.umms.med.umich.edu/) which provides curated RRN data (Stoddard et al., 2015). In order to remove phylogenetic redundancy from the dataset we aggregated mean trait values for all genomes at the species level. The affiliation of genomes at the species level was determined via the GTDB taxonomy (Chaumeil et al., 2020). In the case of genomes that could not be annotated at the species level to the GTDB taxonomy, we considered genomes sharing >94% of their average nucleotide identity as affiliating to the same species (Konstantinidis and Tiedje, 2005). Scripts that were used for determining the here presented genomic traits and the assignment of genomes to species are available via GitHub (https://github.com/sarabeier/genomic.traits) and a trait table is provided in the supplementary material (Supplementary Table S1).

Covariation patterns among the above introduced genomic traits in the multivariate trait space were illustrated via a principal component analysis (PCA, Figure 3). The first two principal components explained > 65% of total variability, while the remaining principal components each contributed < 10% to total variability (Supplementary Figure S3). A random removal of 1, 2, 3 or 4 variables from the PCA demonstrated that the covariation patterns among the remaining genomic traits patterns stayed robust (Supplementary Figure S4). An inspection of individual pairwise correlation strengths (Figure 4) revealed that the absolute spearman rank correlation index rho ranged from 0.04 to 0.92. All correlations were significant due to the large number of included genomes (8,847 species, p-value adjusted via Bonferroni correction for multiple comparisons: Padj < 0.05). Below we will highlight some of the 35 pairwise correlations whose strength or shape displayed via local fitting can be interpreted well in the light of possible physiological constraints.

**Figure 3:**
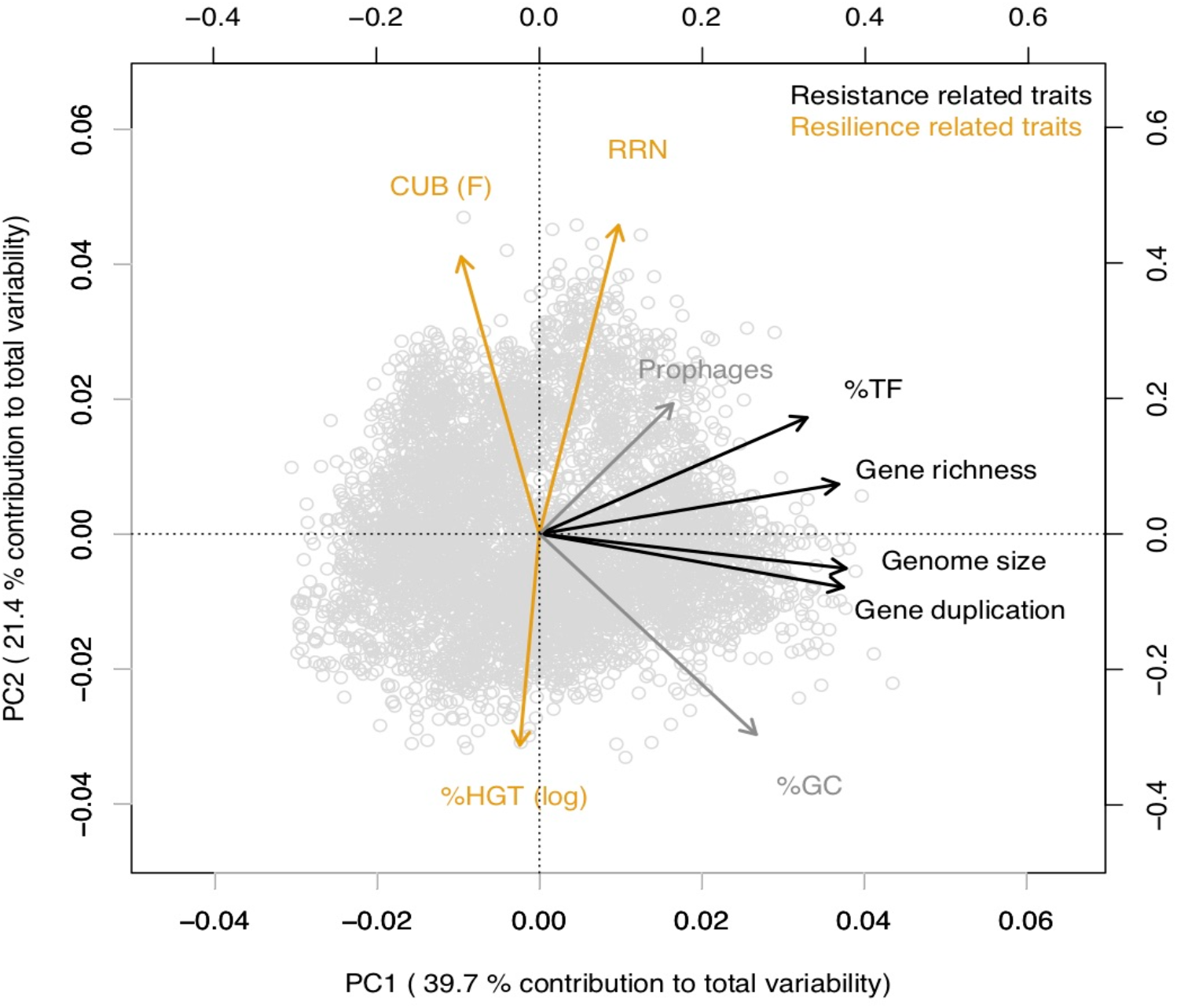
Principal component analysis illustrating covariations among genomic traits from 17,856 sequenced prokaryotic genomes available via the JGI/IMG platform (https://img.igi.doe.gov/) that were averaged at the species level (Supplementary Table S1). 5,823 out 8,847 species shared values for all considered traits and were included in the analyses. The value for %HGT was log(x+0.001) transformed to approximate a normal distribution. We have chosen to display the CUB parameter F (Vieira-Silva and Rocha, 2010) instead of generation time estimations that can be delineated from the CUB, because F is unambiguously defined for all genomes. In contrast, generation time estimations have been suggested to be inaccurate for genomes with large CUB values.

**Figure 4:**
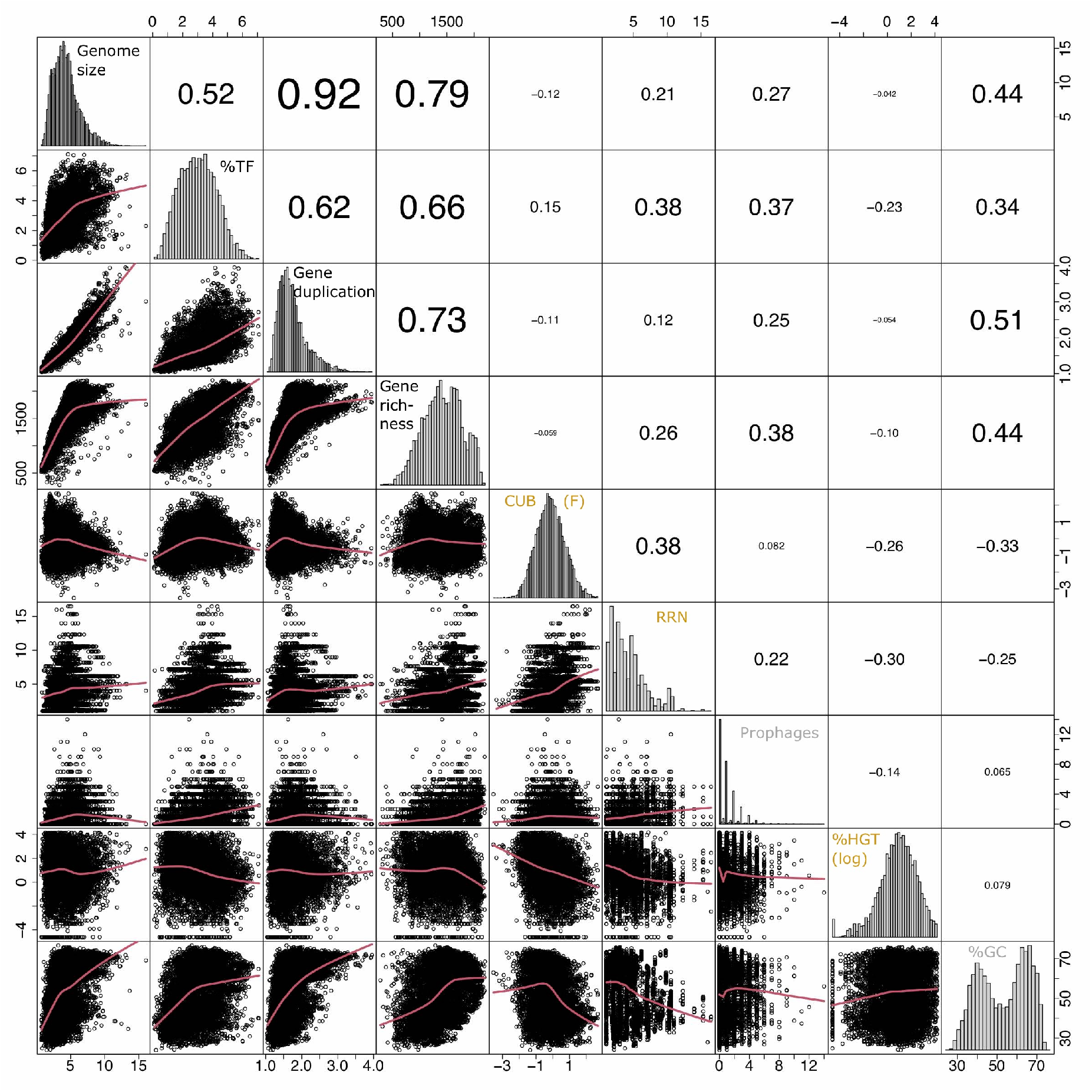
Overview of pairwise correlation patterns among genomic traits from 8,847 species delineated from sequenced prokaryotic genomes available via the JGI/IMG platform (https://img.jgi.doe.gov/) and created via the R command chart.Correlation in the R package PerformanceAnalytics (v2.0.4). The lower panels illustrate pairwise correlation plots fit via loess smoothing statistics with the smoothing parameter f set to 2/3 and the number of robustness iterations set to 3. The value in the upper panels indicate strength of the pairwise correlations (rho, Spearman rank correlation). The diagonal panels illustrate the distribution of the input variables. The CUB was estimated via the variable F as detailed elsewhere (Vieira-Silva and Rocha, 2010). The value for %HGT was log(x+0.001) transformed to approximate normal distribution. The variables RRN and prophages were not transformed as no transformation option improved the fit to normal distribution. The raw values of all data are given in Supplementary Table S1.

Genome size, gene duplication level, gene richness and the %TF aligned approximately along the first principal component (PC1) while covarying positively with each other (Figure 3). An inspection of the pairwise scatter plots revealed a pronounced linear relationship between the genome size and the gene duplication level (Figure 4). In contrast, the increase of gene richness along with genome size rather followed a saturation curve. Apparently, the enrichment of genomes with new genes occurred only until a certain genome size threshold (~ five million base pairs), after which a further genome size increase was mostly due to the duplication of already present genes. A similar saturation pattern was observed for the pairwise correlation of the fraction transcription factors against genome size (Figure 4): a steep positive relationship was apparent approximately up to the genome size until which gene richness increased, while after that threshold a less pronounced increase was observed. Obviously, acquiring new genes requires a stronger enrichment in regulatory genes, than does the duplication of genes. This observation implies that multicopy genes are often, although not explicitly, under the control of the same regulatory operon and can in this case not be expressed alternatively in response to changing conditions. An increasing tolerance to environmental changes due to a genome size increase should therefore, in the case of larger genomes, be primarily due to the dosage effect of replicated genes. The shape of the above reported pairwise relationships accordingly illustrates the physiological mechanisms that link the variables for %TF and genome size depending on whether genome size increases due to the acquisition of new genes or gene duplication. The positive covariation among all four traits underlines their common association with species classifications along the specialist-generalist continuum suggested in literature for genome size and the %TF (Figure 1).

Earlier studies suggested moreover that the % HGT, the CUB and the RRN were either linked to generalist-specialist classifications or correlated positively with genome size (Figure 1, 2), which would imply an alignment of these variables along PC1. Yet, these variables aligned rather along the second principal component (PC2) and were accordingly only weakly correlated to either of the four above described resistance related variables (Figure 3,4). Indeed, a weak correlation between genome size and growth rates that had been observed in several earlier studies (Figure 2, Vieira-Silva and Rocha, 2010; Westoby et al., 2021b) was supported by the likewise weak overall correlation between genomes size and CUB (rho=-0.12) in our analysis.

However, when inspecting the pairwise scatterplots in more details, local fitting suggested a hump-shaped relationship between CUB and genome size, that to our knowledge has not yet been described elsewhere: a positive trendline occurred until a genome size of roughly five million base pairs, after which the direction turned into a negative trendline (Figure 4). Obviously, the observed hump-shaped trendline appeared mainly due to the absence of very small as well as very large genomes with high CUB values. In contrast, genomes with intermediate genome sizes were associated with almost the full range of possible CUB values, resulting in a kind of pyramid-shaped distribution of data points in the pairwise scatterplots. A recent study highlighted that the relationship between CUB values and minimal generation times is inaccurate for prokaryotes with minimal generation times > 5h (Weissman et al., 2021), which complicates the ecological interpretation of CUB values. We therefore want to point out that the observation of prokaryotes with very large or small being exclusively slow growers was not impacted by these inaccuracies in the relationship between CUB values and generation times (Figure 5).

**Figure 5:**
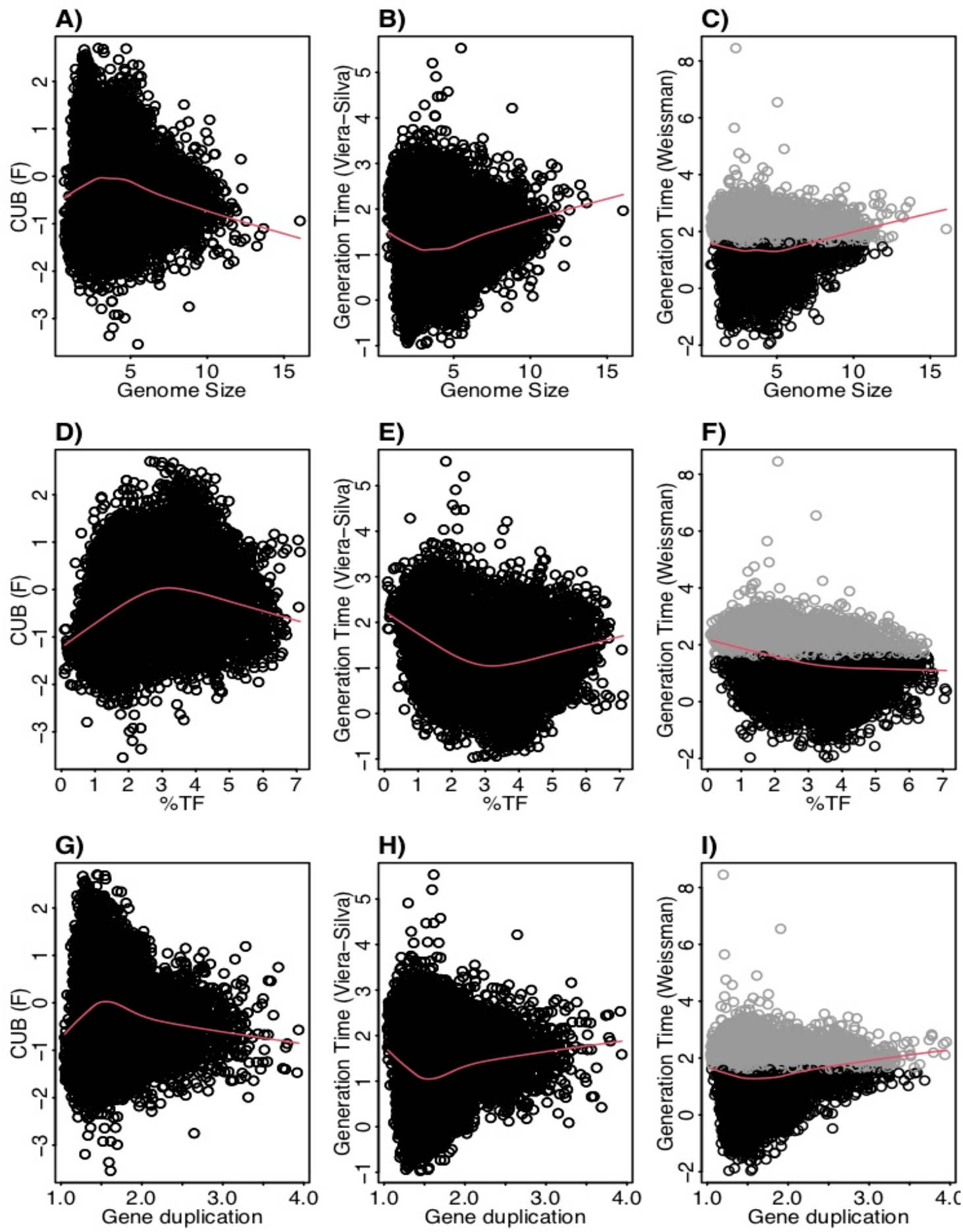
Scatterplots displaying pairwise relationships between the codon usage bias (F, Vieira-Silva and Rocha, 2010) the generation time (log-transformed) estimated as detailed in Vieira-Silva et al. (2010) and the generation time (log-transformed) estimated using the R package gRodon (v0.0.0.9000, Weissman et al., 2021) against genomes size, %TF and gene duplication. Genomes with low codon usage bias resulting in predicted generations times (gRodon) exceeding 5 h (highlighted in gray) represent species with generations times >5 h, while the exact value is inaccurate. The absence of genomes with elevated generation times at the extremes of the value ranges for the displayed resistance related traits is accordingly independent from these inaccuracies.

Physiological reasons for a non-monotone relationship between growth rate and genome size could be due to opposing mechanisms that prevail under different genome sizes ranges: on the one hand, it had been proposed that an increasing proportion of genes involved in metabolic pathways, that was observed along with increasing genome sizes (Konstantinidis and Tiedje, 2004) causes higher metabolic rates. Consequently, an increased availability of energy supply via ATP should lead to enhanced growth rates (DeLong et al., 2010). In this scenario, the growth rate of very small genomes would be limited by the available energy. On the other hand, it has been argued that cells with small genomes feature higher growth rates than cells with large genomes, because they can initiate a new replication cycle before the previous rounds have been finished (Vieira-Silva and Rocha, 2010). Furthermore, the proportion of genes encoding the translation of mRNA into proteins, DNA replication or cell division is reduced in large genomes (Konstantinidis and Tiedje, 2004), which could lead to reduced growth rates. We propose that these latter two physiological constraints limit the growth rates of species with very large genomes. It has been discussed earlier that genome size reduction in prokaryotes can on the one hand be induced via genetic drift, a mechanism that should affect particularly intracellular parasites with small effective population sizes. Conversely, genome streamlining due to adaptive selection occur typically in response to nutrient limitation in aquatic systems (Giovannoni et al., 2014). Still, the above highlighted physiological consequences of genome size reduction should apply to all small genomes. Indeed, both, intracellular parasites as well free living oligotrophic organisms with small genome sizes are typically characterized as slow growing organisms that are sensitive to environmental change (Couturier and Rocha, 2006; Joseph and Goebel, 2007; Parter et al., 2007; Giovannoni et al., 2014). We argue based on these considerations that genome size itself causes the above described physiological constraints and characteristics, independent from the evolutionary forces selecting for prokaryotes with enlarged or reduced genome size.

A trend for hump-shaped relationships and / or pyramid-shaped distribution of data points in the pairwise scatterplots was not only visible if plotting genome sizes against CUB values: several other pair-wise comparisons between traits associated with the specialist-generalist continuum (genome size, gene duplication level and the fraction of transcription factors) versus and traits that covaried positively with CUB (RRN, number of prophages) displayed similar profiles (Figure 4).

The positive covariation between the number of prophages with the RRN and, to a certain extent, with CUB and, accordingly, with maximum growth rate has already been outlined earlier (Touchon et al., 2016). It has been argued that a higher number of prophages in potentially fast growing bacteria is a consequence of their opportunist life style that provides more variable growth states and resources for the production of virions. This in turn was suggested to favor lysogeny and therefore the presence of prophages (Touchon et al., 2016). However, the number of prophages was placed in-between the PCA axes PC1 and PC2 and also covaried positively with %TF and gene richness. This seems reasonable as the integration of prophages into the genome adds more genes to the genome. The lateral transfer of genes via prophages is one of several mechanisms leading to HGT. It is therefore remarkable that we found a negative correlation between the number of prophages with the %HGT (Figure 3). We conclude from this observation that the overlap between the detected HGT events and prophages was low. This is possibly due to a high host-specificity of bacteriophages, which would lead to gene transfer only between closely related strains. We assume that the phylogeny-based method that was applied to detect the %HGT in the JGI genome statistics and that identifies genes with phylogenetic foreign origin in genomes, does not resolve well genes acquired from close relatives (Markowitz et al., 2010).

The %HGT covaried not only negatively with the number of prophages, but also with CUB (Figure 3). Such negative covariation has to our knowledge not yet been reported. However, the discovery that the %HGT events in genomes is connected to their optimal growth temperature (Gophna et al., 2015) may be linked to the above mentioned negative covariation, as species with high growth optimum tend to have comparably high maximal growth rate (i.e. low minimal generation time; Vieira-Silva and Rocha, 2010). A high degree of sub-species genome plasticity in the typically slow growing members of the SAR11 clade (Ward et al., 2017) is in agreement with this observation. It indeed seems reasonable that a fine tuned genetic replication machinery is necessary for achieving high maximal growth rate, which may easily be impeded by a high fraction of genes of foreign origin. Our data therefore point to a possible tradeoff between high growth rates and the ability of genomes to stably integrate foreign DNA from different phylogenetic origin. It needs though to be considered that the phylogeny-based methods to detect %HGT events may overestimate HGT events for genomes with few closely related genomes that belong to the same phylogenetic group. The above suggested tradeoff should therefore be confirmed in other datasets, possibly in combination with other approaches to detected HGT events.

The GC content did not exhibit a pronounced covariation with any other genomic trait. However, a certain level of covariation could be detected with genome size and the gene duplication level or with CUB and the fraction of horizontally transferred genes (Figure 3). The limited covariation of the GC content with any other trait could be due to the complex selection forces, including on the one hand nutrient availability and on the other hand exposure to heat or desiccation stress that in combination drive the GC content evolution (Hellweger et al., 2018; Chen et al., 2020).

## 4. Synthesis

Based on the covariation patterns observed here in combination with findings from earlier studies, we propose that the %TF, genome size, gene richness and gene duplication represent resistant related traits and align approximately with the first principal component of the PCA (Figure 3). In contrast, traits including CUB, RRN and %HGT that according to earlier studies or in agreement with our analyses exhibited covariations with growth rates or lag phases and that hence could be considered as resilience related traits aligned rather with the second principal component of the PCA (Figure 3).

We did not assign %GC to resistance or resilience related traits as it was positioned in-between PC1 and PC2 and did not align closely to any of the other traits. Although a high GC content can protect cells from heat or desiccation stress resistance, it can furthermore not be considered as a general resistance trait enabling for instance tolerance against changes in the resource type supply. Likewise, we did not assign the number of prophages to either resistance nor resilience related traits based on its position in-between PC1 and PC2. Apart from an earlier suggested link between an elevated number of prophages with high growth rates and an opportunistic lifestyle (Touchon et al., 2016) an increasing number of prophages will also increase the genome size and other resistance related traits.

The not fully congruent overlap between the genomic traits that we assigned to either resistance or resilient related traits suggests that these traits cover different aspects of resistance and resilience, such as increased growth rates versus shorter lag phases in the case of resilience related traits.

The roughly orthogonal position of several resistance and resilience traits in prokaryotes across whole range genome sizes reflects the results of some previous studies, which suggested that genome size and growth rate are not related (Vieira-Silva and Rocha, 2010; Westoby et al., 2021b). However, the analyses presented here were based on a database that exceeded those used in these former studies by one to two orders of magnitude. An inspection of pairwise trait correlations indicated the presence of non-monotone relationships between several resistance and resilience related genomic traits that to our knowledge were not yet reported in earlier studies: for instance, genomes with a size up to roughly four million base pairs featured according to a local fitting approach a positive relationship with CUB, while after approximately five million base pairs it turned to a negative relationship (Figure 4, 5). Furthermore, genomes up to this threshold continued to increase due to a combination of newly acquired genes and gene duplication events, while larger genomes increased rather due to gene duplication events (Figure 4). Last but not least aquatic environments are typically characterized by genomes smaller than four million base pairs, while soil environments usually harbor genomes larger than five million base pairs (Giovannoni et al., 2014). Correlation analyses of partial datasets exhibited in agreement with the trendlines obtained by local fitting significant positive correlations between several resistance versus resilience related traits, if considering genomes up to a size of four million base pairs. The opposite was true in most cases if considering genomes larger than five million base pairs (Table 1). Results from these partial correlation analyses (Table 1) highlight that non-monotone relationships are sensitive to the range of data points included. We argue that non-monotone relationships are the reason for contradicting findings e.g. the relationship between genome size and growth rates, that have been described as either positively related (Freilich et al., 2009; DeLong et al., 2010) or as largely unrelated dimensions (Vieira-Silva and Rocha, 2010; Westoby et al., 2021b). Furthermore, the previously reported superlinear positive correlation between genome size and HGT events (Cordero and Hogeweg, 2009b) or the assignment of generalist species to high %HGT, large CUBs or increased growth rates (Figure 2), which was not supported by our findings, could be due to the specific set of genomes that were used in the respective analyses.

**Table 1:**
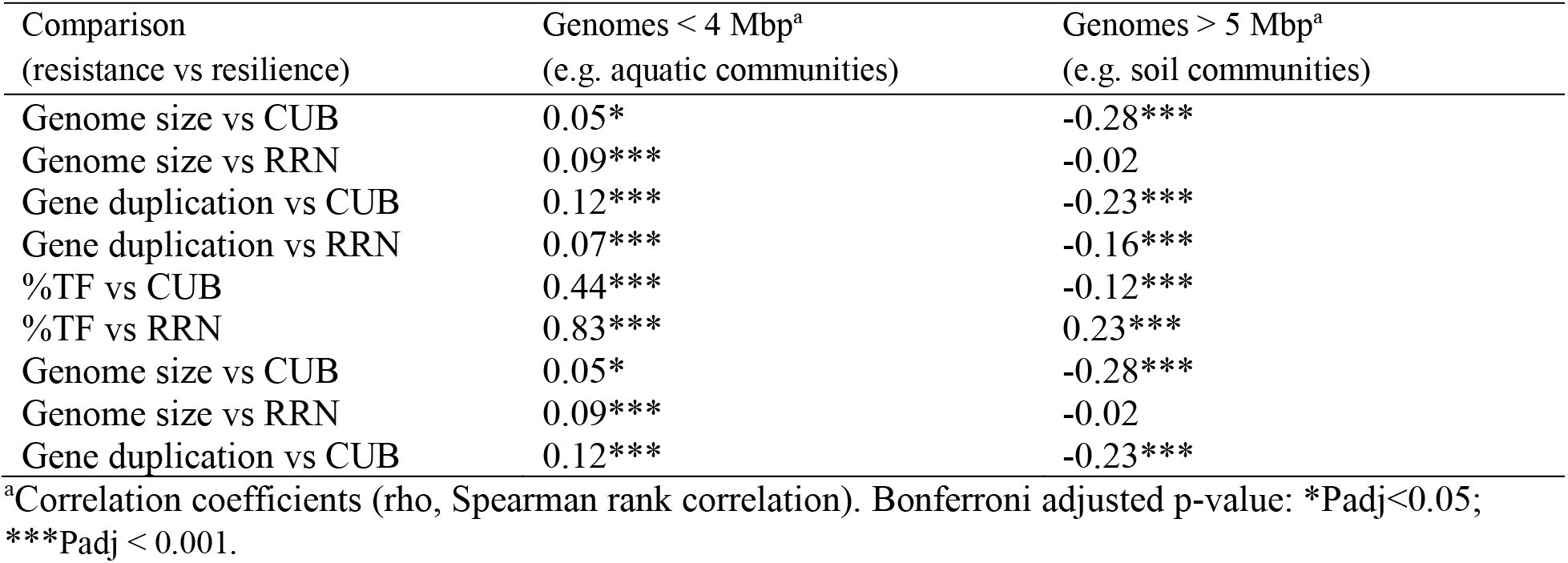
Correlation coefficients between selected resistance and resilience related traits for species with small and large genomes

**Table 2:** Phylogenetic signals of genomic traits

| Genomic trait | K<sup>a</sup> | P (K)<sup>a</sup> | Lambda<sup>b</sup> | P (lambda)<sup>b</sup> | Max PD<sup>c</sup> |
| --- | --- | --- | --- | --- | --- |
| Genome size (JGI/IMG)<sup>d</sup> | $5.89 \times 10^{-4}$ | <0.001 | 0.9973 | <0.001 | 1.5 |
| %TF (JGI/IMG) | $3.10 \times 10^{-4}$ | <0.001 | 0.9905 | <0.001 | 1.5 |
| Gene duplication (JGI/IMG) | $4.89 \times 10^{-4}$ | <0.001 | 0.9976 | <0.001 | 1.5 |
| Gene richness (JGI/IMG) | $5.52 \times 10^{-4}$ | <0.001 | 0.9892 | <0.001 | 2.5 |
| CUB (F, JGI/IMG) | $1.24 \times 10^{-3}$ | <0.001 | 0.9974 | <0.001 | 1.5 |
| Generation time (gRodon, JGI/IMG) | $4.66 \times 10^{-4}$ | <0.001 | 0.9935 | <0.001 | 2.0 |
| RRN (rrnDB)<sup>d</sup> | $1.20 \times 10^{-6}$ | <0.001 | 0.9624 | <0.001 | 1.5 |
| Prophages (JGI/IMG) | $1.33 \times 10^{-5}$ | <0.001 | 0.6403 | <0.001 | 0.5 |
| %HGT (JGI/IMG) | $4.22 \times 10^{-5}$ | <0.001 | 0.8897 | <0.001 | 1.5 |
| %GC (JGI/IMG) | $2.33 \times 10^{-2}$ | <0.001 | 0.9999 | <0.001 | 1.5 |

<sup>a</sup>Blomberg's K statistics.
<sup>b</sup>Pagel's Lambda statistics.
<sup>d</sup> In the specific case of RRN we provide phylogenetic signal estimated based on trait values from the rrnDB, while for all other traits JGI/IMG trait values as reported in the Supplementary Table S1 were used (%HGT was $\log(x+0.001)$ transformed and Generation time was $\log(x)$ transformed). We inferred the RRN rrnDB phylogenetic signal from a phylogenetic tree calculated using rrnDB 16s rRNA gene sequences (FastTree 2, Price et al., 2010) and using the trait values available via the rrnDB (https://rrnDB.umms.med.umich.edu/static/download/, rrnDB-5.7). The phylogeny for all JGI/IMG values was inferred from the pro\_ref.tre phylogenetic tree, which is used as backbone phylogeny of the PICRUST2 software. All analyses to compute phylogenetic signals were performed in R (R Core Team, 2021) and a script is available on GitHub (https://github.com/sarabeier/genomic.traits).

Noticeably, not all partial pairwise correlations were strong or resulted in a differential correlation profile for larger and smaller genomes (Table 1). Still, we believe that as a consequence of the inconsistent correlation patterns reported here, positive relationships between resistance and resilience are more likely to occur in aquatic habitats: a positive covariation of genomic traits that we here classified as resistance and resilience related traits had been recently observed in an aquatic fertilization experiment (Okie et al., 2020). Instead, tradeoffs between resistance and resilience should be more likely in soil habitats. Indeed, the above outlined tradeoff between functional resistance and resilience has to our knowledge mainly been observed in soil environments (e.g., Garcia et al., 2020; Piton et al., 2021). This is in agreement with the negative correlation between several resistance and resilience related traits particularly among larger genomes. Habitat specific PCA patterns with genomes originating from soils, aquatic habitats or from the digestive tract furthermore supported our findings: the position of resistance versus resilience related traits shifted with increasing size of the included genomes from positive towards negative covariations along the first principal component (Figure 6). The here observed shifts may be even more extreme in natural habitats, as a considerable part of genomes in the reference database originated from cultured strains. Particularly in aquatic habitats these are likely biased towards larger genomes while oligotroph large genomes in soil habitats might be underrepresented, thereby resulting in genome size distributions that are not typical for these habitat types.

**Figure 6:**
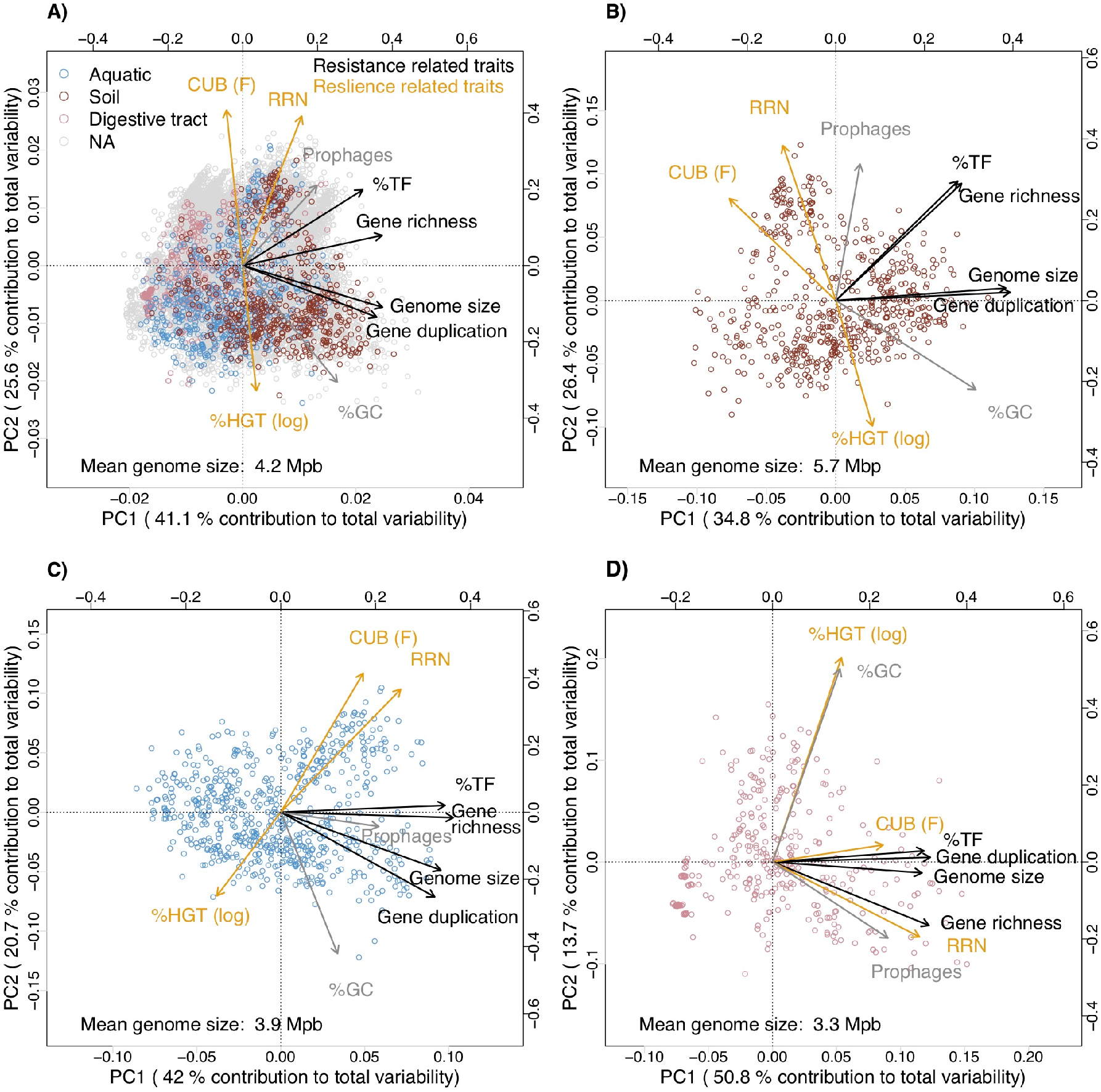
Principal component analyses illustrating covariations among genomic traits from 17,856 JGI/IMG prokaryotic genomes dependence on the habitat type. A) All genomes. B) Genomes originating from soil habitats. C) Genomes originating from aquatic habitats. D) Genomes originating from the digestive tract. In order to display three habitat types from which individual genomes originated, we did not aggregate the genomes at the species level. The three habitat types were determined via text search from the habitat information available via the JGI/IMG database (Supplementary Table S1; ignore.case=TRUE): all genomes containing the strings ‘soil’ or ‘rhizosphere’ in the habitat description where classified as originating from soil habitats; all genomes containing the strings ‘aquatic’ or ‘marine’ or ‘water’ where classified as originating from aquatic habitats; all genomes containing the strings ‘oral’ or ‘stomach’ or ‘gut’ or ‘intestinal’ or ‘feces’ were classified as originating from the intestinal tract. The remaining genomes were not further classified.

**Figure 7:**
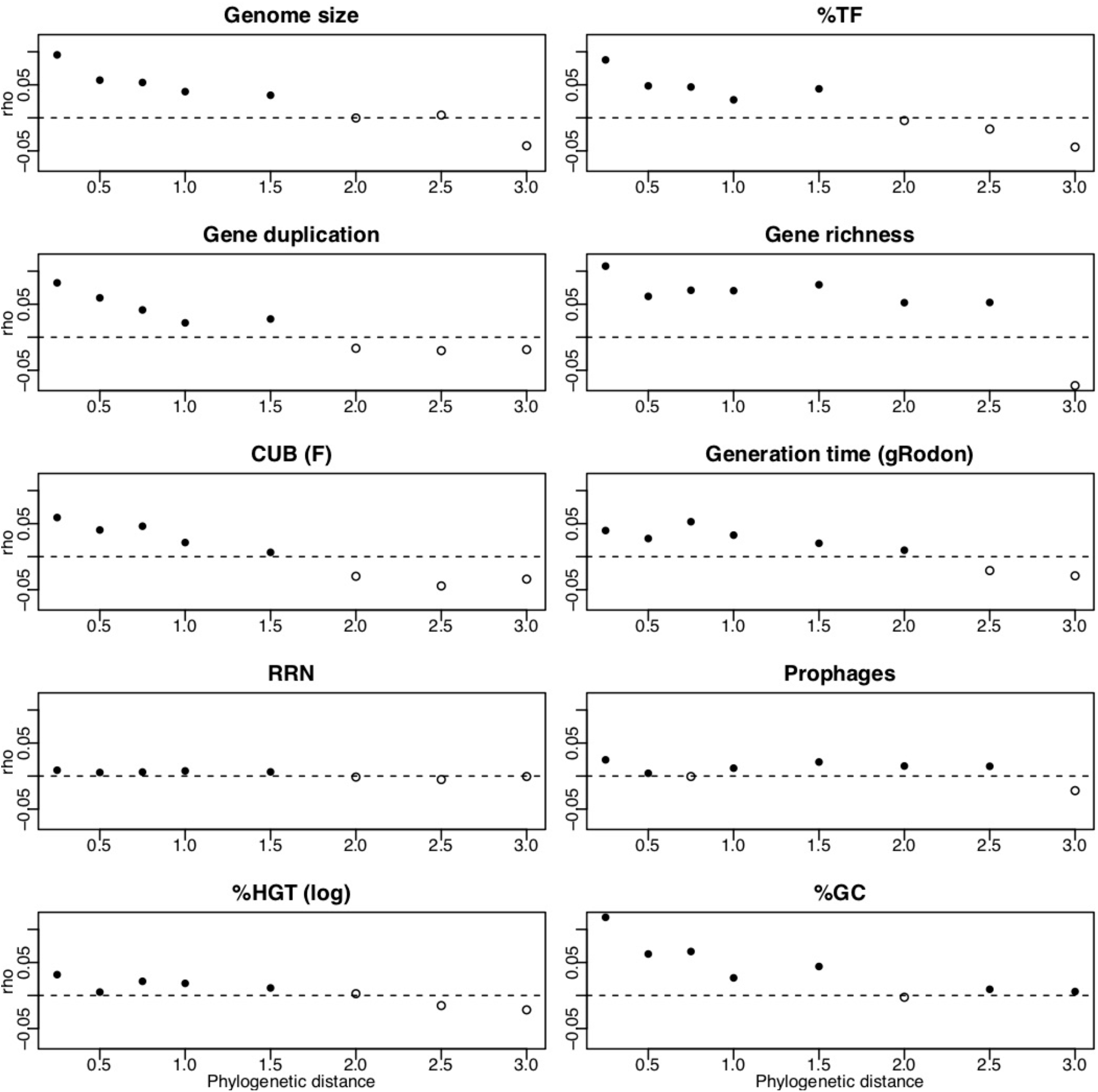
Results from Mantel correlograms. Filled data points indicate positive significant correlations (Bonferroni adjusted p-values Padj < 0.1) of pairwise distances of traits values against pairwise phylogenetic distances among the reference genomes. The Mantel correlograms were computed for 10,000 randomly selected genomes and with 200 permutations. In the specific case of RRN we computed the Mantel correlogram based on trait values of the rrnDB, while for all other traits JGI/IMG trait values as reported in the Supplementary Table S1 were used (%HGT was log(x+0.001) transformed and Generation time was log(x) transformed). We inferred the RRN rrnDB phylogenetic signal from a phylogenetic tree calculated using rrndbDB 16s rRNA gene sequences (FastTree 2, Price et al., 2010) and using the trait values available via the rrnDB (https://rrndb.umms.med.umich.edu/static/download/, rrnDB-5.7). The phylogeny for all JGI/IMG values was inferred using the pro_ref.tre phylogenetic tree, which is used as backbone phylogeny of the PICRUSt2 software. All analyses to compute Mantel correlograms were performed in R (R Core Team, 2021). Scripts with the code for all analyses described in this figure legend are available on GitHub (https://github.com/sarabeier/genomic.traits).

Literature suggests consistently that increasing resource availability leads to the selection of fast growing opportunists (i.e. r-strategists) with decreased resource usage efficiency (Stevenson and Schmidt, 2004; Fierer et al., 2007; Roller et al., 2016). Prokaryotes with high resource usage efficiency and simultaneously low maximal growth rates are usually observed in oligotroph environments with the possible exception of obligate intracellular living bacteria: as mentioned above is the evolution of these organism largely affected by genetic drift and they typically live in nutrient rich environments, but tend to feature low growth rates (Couturier and Rocha, 2006; Joseph and Goebel, 2007). We suggest copiotrophic-oligotrophic classifications, at least in free living communities with large effective population sizes that are little impacted by genetic drift, to be aligned with the dimension of resilience and resource usage efficiency. This implies that classifications of free living communities along the copiotrophic-oligotrophic axis are analogously to the resistance axis decoupled from the dimension of resistance related classifications if considering species along the whole range of genome sizes. However, contrasting relationships can occur for instance in aquatic versus soil habitats. In agreement with these theoretical considerations, previous studies actually reported the selection of larger genomes after nutrient addition in aquatic habitats and the opposite in soil habitats (Figure 2). In line with this, a recent study indicated based on habitat dependent covariation patterns between genome size and %GC that genome reduction in soil habitats may not be driven by known mechanisms, such as streamlining due to nutrient limitation (Chuckran et al., 2021).

Importantly, the proposed decoupling of resistance versus resilience and resource usage efficiency dimensions entails that a consistent assignment of traits to the CSR or YAS schemes with validity for all prokaryotes may not be possible: in environments inhabited by prokaryotes with small genomes, such as aquatic habitats a nutrient rich and frequently disturbed habitat should select species with increased CUB and comparably large genomes, while the same scenario in soil environments should instead rather select species with increased CUB, but comparably small genomes. Genome size dependent trait-trait covariation patters might furthermore be the reason for conflicting assignments of genome size either with the C (Fierer et al., 2007) or R category (Krause et al., 2014) and likewise RRN or CUB/growth rate with either the R and S categories (Fierer et al., 2007) or with the C category (Krause et al., 2014) of the CSR schema.

We want to emphasize that the classification of individual species via their genomic traits into life history categories suffers from inaccuracies. This is on the one hand due to the fact that different evolutionary mechanisms and selective forces in combination lead to the selection of genomic trait values, which causes noisy correlations between trait values and the functional characteristics of a species. On the other hand, the assignment of species for instance along the specialist-generalist gradient is highly context-dependent and a resource specialist might be at the same time a salinity generalist (Bell and Bell, 2021). As a consequence, it is not possible to unambiguously characterize prokaryote species along the generalist-specialist continuum or predict their phenotypic response to a specific environmental change based on a simple genomic trait as for instance their genome size. Still, the probability that a species will be resistant against a specific environmental change increases with its genome size. Similarly, while it is not possible to predict from a high CUB that the corresponding species will actually grow fast in a given environment, a high CUB increases the probability of this species to exhibit high growth rates in this environment. Accordingly, although the outlined genomic traits are imprecise in predicting the phenotypic characteristics of individual species in a given environment, they affect its likelihood to be tolerant or to grow fast in this environment. While probabilities do not allow the prediction of a single event, their predictive power increases with the number of considered events. We therefore suggest that resistance or resilience of individual species in a given situation should preferably be evaluated via the regulation of RNA markers in response to a specific environmental change, as detailed in a recent study (Rain-Franco et al., 2021). We however claim that the predictability of functional consequences from genomic traits increases if simultaneously applied for multiple species in a community and genomic traits values are scaled-up to evaluate their distribution at the community level (BOX 2).

## 5. Conclusions

Recent publications claimed that trait dimensions that are apparent among heterotroph prokaryotes are due to different physiological constraints and tradeoffs not directly comparable to those of autotroph plants (Malik et al., 2020; Westoby et al., 2021a). Based on our analyses we suggest that physiological constraints and tradeoffs differ even within the microbial cosmos, which precludes a globally consistent assignment of microbial traits in agreement with the CSR or YAS frameworks. In contrast, sorting microbial traits within a resistance/resilience framework increased consistency between trait-trait covariations and earlier reported findings due to variable tradeoffs between resistance and resilience related traits in dependence of the genome size range. It has been argued that varying resistance/resilience relationship have consequences for the stability of communities: In aquatic systems, where our analyses suggest a high likelihood for positive relationship between resistance and resilience levels, disturbances under oligotroph conditions should lead according to ecological theory to a species loss because both resistance and resilience are simultaneously low (Nimmo et al., 2015). Disturbances under high resource availability should instead induce a gain of species if resistance and residence levels are simultaneously high. In contrast, theory suggests a higher degree of stability in response to disturbances in communities featuring a resistance/resilience tradeoff (i.e. soil communities) because these communities are either resistant or resilient (Nimmo et al., 2015).

Even though specific CSR or YAS trait attributions may not generally be applicable for all prokaryotes, they should be applicable with adapted trait associations for communities harboring species within certain ranges of genome sizes, such as aquatic or soil communities. We argue that, beside potential differences between heterotrophs and autotrophs outlined elsewhere (Malik et al., 2020; Westoby et al., 2021a), disturbances and productivity gradients are, analogous to plants, the main drivers for microbial community dynamics. To understand the ecology of microbes and make predictions about their dynamics it is consequently essential to combine different trait dimensions, concerning their response to disturbances and nutrient availability. We emphasize the need to expose prokaryote communities from different habitats to experimentally crossed disturbance and productivity gradients using full factorial designs and examine genomic trait distributions and diversity patterns. This should ideally be done in combination with functional resistance and resilience measurements to validate the links between genomic traits and community-level functional characteristics of prokaryotes outlined in this study. Such experimental designs will enable to empirically underpin the here presented predictions concerning the assembly of genomic traits and species richness under different scenarios and their relevance for community functioning. The trait table used in this study may hereby serve to extrapolate genomic trait distributions in prokaryotic communities based on taxonomic marker genes as outlined in BOX2.

### BOX 2: Evaluation of genomic trait distributions from community sequence data

The shape of trait distributions in communities represented by measures such as the community weighted mean (CWM), but also the community weighted variance, skewness or kurtosis belong to the key drivers of community functioning and assembly (Enquist et al., 2015). While there are practical constraints to measure the distribution of physiological traits in microbial communities, genomic trait distributions can be extracted from community sequencing data. For instance, the CWM of the GC contents or genome sizes in microbial communities can be determined directly from the sequenced reads of shot gun metagenomes. In the latter case this can be done by relating the number of reads coding for single copy housekeeping genes to the number of total reads (Nayfach and Pollard, 2015). The same is true for the RRN, as the number of reads coding for 16s rRNA genes can be directly identified from metagenome reads (Kopylova et al., 2012) and related to the number of reads coding for single copy housekeeping genes (Biers et al., 2009). However, while the typically highly conserved housekeeping genes can be identified with high precision from short sequence reads the functional annotation of less conserved genes from short reads lacks accuracy. Longer sequences are therefore necessary for the Hidden Markov Models based annotation of genes encoding transcription factors as well as for the CUB estimation (Vieira-Silva and Rocha, 2010). In order to estimate the gene richness or gene duplication level within genomes as well as the number of prophages, an access to (nearly) full genome sequences is necessary. Longer sequences and even assembled genome sequences from shotgun metagenome data can be obtained via assembly and genome binning approaches. However, both methods (particularly genome binning) are biased towards more abundant sequences and genomes, while some life history traits may prevail in the rare biosphere (Vergin et al., 2013). Accordingly, may genomic traits representing the life histories of species that are rare in a community be underestimated if assembled or binned metagenome data were used for trait detection.

Another option to assess the distribution of genomic traits is to infer genomic traits from taxonomic marker genes of species in a community based on sequenced reference genomes of close relatives (Cébron et al., 2021; Romillac and Santorufo, 2021). This procedure is possible for all traits featuring a sufficiently strong phylogenetic signal (BOX3) and has been applied to determine genome sizes (Barberan et al., 2014) or the CUB (Weissman et al., 2021) of microbial communities based on 16s rRNA gene sequence data. Beside avoiding the possible biases outlined above, this strategy would allow to not only determine CWS, but also the other moments of trait distributions and thereby enable a more thorough evaluation of trait distributions in microbial communities.

The PICRUSt2 software (Douglas et al., 2020) that had been designed to extrapolate the genomic content of uncultured prokaryotes from closely related genomes via taxonomic marker genes can be analogously used to extrapolate the genomic traits outlined here. Our trait table (Table S1) can be applied to predict genomic traits for each species (OTU/ASV) in a sequenced community via the hidden state prediction tool (Louca and Doebeli, 2018) integrated into the PICRUSt2 software and using the default PICRUSt2 species reference database (PICRUSt does not accept trait values outside the range of 0-1000 nor values that have more than two decimal digits, accordingly some of the given trait values need to be re-ranged and rounded). Douglas and colleagues present averaged information of several genomes, if these contained identical 16s rRNA genes (Table S1: picrust.ID = *-cluster), while the trait data presented in Table S1 (with exception of the RRN) refer exclusively to the genomes indicated in column IMG.Genome.ID. However, due to the highly significant phylogenetic signals detected for the presented genomic traits (Table 2), we believe that the variability of these traits among close relatives sharing the same 16s rRNA gene should be limited. In the particular case of RRN and as a consequence of putatively biased RRN values of the JGI/IMG database (Supplementary Figures S1/S2), we propose to use values given in the rrnDB database (Stoddard et al., 2015) in combination with the respective phylogeny to predict RRN values from amplicon sequence data.

### BOX 3: Phylogenetic signals of genomic traits

All above evaluated genomic traits featured overall significant phylogenetic signals (Table 2). In agreement with the phylogenetic signals, Mantel correlograms illustrated significant positive correlations between phylogenetic and trait value distances at least until a phylogenetic distance of 0.25 (Table 2, Figure 7). The PICRUSt2 software (Douglas et al., 2020) provides phylogenetic distances to the nearest relative in the reference database for each OTU/ASV in a sequenced query community (NSTI values) and a default cutoff level of NSTI<0.2 is used to exclude predictions for OTUs/ASV that are not represented by close relatives in the PICRUSt database. NSTI values can be used in combination with the here reported results from the mantel correlograms to evaluate the reliability for PICRUSt-based trait predictions: for instance the community average NSTI value (weighted mean) in a soil community computed with the PICRUSt1 software was 0.17 (Langille et al., 2013).

Due to possible biases of RRN values given in the JGI database (Figure S1,S2) we estimated the phylogenetic signal for RRN based on original entries of the rrnDB in combination with the corresponding rrnDB phylogeny. We assume that the comparably low phylogenetic signals detected for RRN (Table 2, Figure 7) may not be a purely biological signal, but could be due to inaccuracies of this parameter that, although not as pronounced as in the JGI database, seemed to be also inherent in the rrnDB database (Figure S2).

One may wonder why genomic traits, such as prophages or % HGT at all exhibit phylogenetic signals, although individual events contributing to these traits are not inherited vertically. However, while the presence of a specific prophage or HGT gene in a genome should indeed not have a phylogenetic signal, it has been argued as outlined above that the characteristic of a genome to host multiple prophages or HGT derived genes is linked to the life-history of the corresponding organism. In agreement with our observation (Table 2), life history traits featured comparably strong phylogenetic signals in an earlier study (Blomberg et al., 2003). Still, in the case of prophages and % HGT it seems biologically meaningful that detected phylogenetic signals were low compared those of other traits (with the exception of RRN).

## 6. Data availability and availability of detailed method descriptions

Sequence data for all genomes included in this study are available via the JGI/IMG platform (https://img.jgi.doe.gov/). The scripts used for bioinformatic data processing are available via GitHub (https://github.com/sarabeier/genomic.traits).

## 7. Acknowledgements

We thank Christiane Hassenrück for providing code to extract raw read data for genomes stored in the IMG/JGI database. We also Sara Vieira-Silva for her advices to setup scripts for the estimation of codon usage biases and the associated generation times from genome data. The study was supported by a grant from the German Science Foundation (DFG) awarded to SB (BE 5937/2-1). Bioinformatic analyses were supported by the BMBF-funded de.NBI Cloud within the German Network for Bioinformatics Infrastructure (de.NBI) (031A537B, 031A533A, 031A538A, 031A533B, 031A535A, 031A537C, 031A534A, 031A532B).

## Supplementary Material

Figure S1: A) Correlation of RRN data from the JGI/IMG database (RRN_IMG) against RRN values that were extrapolated from genome level RRN values available via the rrnDB database indicate that a bias in the JGI/IMG values is mainly due to RRN underestimation for entries with high rrnDB RRN values. B) The standard deviation (sd) of within genus (NCBI taxonomy) RRN values for 666 genera of the rrnDB database with more than one entry per genus indicate values of sd < 1 in the large majority of genera.

Figure S2: The JGI database provides information about NCBI accession, bioproject or biosample accession numbers, but no direct link to the raw read data. While for the majority of JGI entries listed in Table S1 it was not possible to identify runids by command-line based matches, we could extract raw read data for some of the entries. We applied the MicrobeCensus software to estimate the genome size as well as the number of sequenced genome equivalents based on the number of reads coding for single copy housekeeping genes relative to the number of total reads (Nayfach and Pollard, 2015). We excluded genomes with <2,000,000 sequenced reads and < 50 sequenced genome equivalents. We further applied the SortMeRNA software (Kopylova et al., 2012) to identify reads coding for the 16s rRNA and used the output in combination with the indicated number of sequenced genome equivalents to estimate the number of RRN per genome. A) Pearson correlation between the read based RRN estimate and the JGI RRN estimate. B) Pearson correlation between the read based RRN estimate and the rrnDB RRN estimate as given in Table S1. C) Pearson correlation between the read based genome size estimate and the genome size given in the JGI database.

While both the read-based and assembly-based estimation of RRN or genome size may suffer from biases, these biases are different and independent from each other. We therefore interpreted the strength of correlation between assembly-based and read-based estimates for RRN and genome size as measure for the quality of values provided in the rrnDB and JGI/IMG databases. The strongly improved correlation of read based RRN estimates against the rrnDB derived RRN values compared to a correlation against the JGI values, indicates that the presented rrnDB derived RRNs are exposed to reduced biases. However, the even better correlation between the genome size estimates in combination with low phylogenetic signals of RRN (Table 2, Figure 7) suggests that also the rrnDB derived RRN are likely still less accurate that are assembly-based genome estimates from the JGI/IMG database.

Figure S3: Percent contribution to variability of principal components PC1 to PC9.

Figure S4: The random removal of 1,2,3 or 4 variable from the principal component analyses presented in Figure 3 demonstrates that the spatial patterns of resistance versus resilience related genomic traits are robust against the removal of individual variables from the dataset.

Table S1: Overview table for 17,856 genomes available via the JGI/IMG platform (https://img.jgi.doe.gov/, downloaded in August 2021), which are integrated in the reference database of the PICRUSt2 software (v2.1.2-b, Douglas et al., 2020). Some genomes present in the original PICRUSt2 reference database had meanwhile been replaced or removed. In the case of replaced genomes we indicate the new IMG genome ID in the column IMG.Genome.ID, while in the column PICRUSt.ID the former IMG genome ID is given. Douglas and colleagues furthermore present averaged information of several genomes, if these contained identical 16s rRNA genes (picrust.ID = *-cluster). However, the trait data presented in Table S1 refer exclusively to the genomes indicated in column IMG.Genome.ID. All traits values were either directly extracted from the JGI/IMG database or computed with scripts that are available on GitHub (https://github.com/sarabeier/genomic.traits). RRN_IMG and RRN_rrnDB represent RRN values given by the JGI/IMG database and values extrapolated at the genus level (NCBI taxonomy) from the rrnDB, respectively. It was not possible to extract the information for all traits from all genomes and missing trait values are indicated by NA. Strains that could not be unambiguously binned at the species level via the GTDB taxonomy were classified via the FastANI software (Jain et al., 2018) into species level bins using an average nucleotide identity >94% (column FastANI_species). Information of all remaining columns is available via the JGI/IMG database.

Table S2: Read based estimation for RRN and genome size of JGI entries with >2,000,000 sequenced reads and >50 sequenced genome equivalents.

## Notes

https://github.com/sarabeier/genomic.traits

https://img.jgi.doe.gov/

https://rrndb.umms.med.umich.edu/

### Competing Interest Statement

The authors have declared no competing interest.

### Summary of Updates

*RRN values of genomes from the JGI/IMG database used in this study were extrapolated from entries of the rrnDB database. This was done because analyses indicated that RRN values given in the JGI/IMG database often underestimate the true RRN. *In order to reduce phylogenetic redundancy of the dataset, we aggregated genomes at the species level (GTDB taxonomy). *Habitat specific PCA plot were added.

