## Supplementary Figures for "Trait-trait relationships and tradeoffs vary with genome size in prokaryotes"

Figure S1

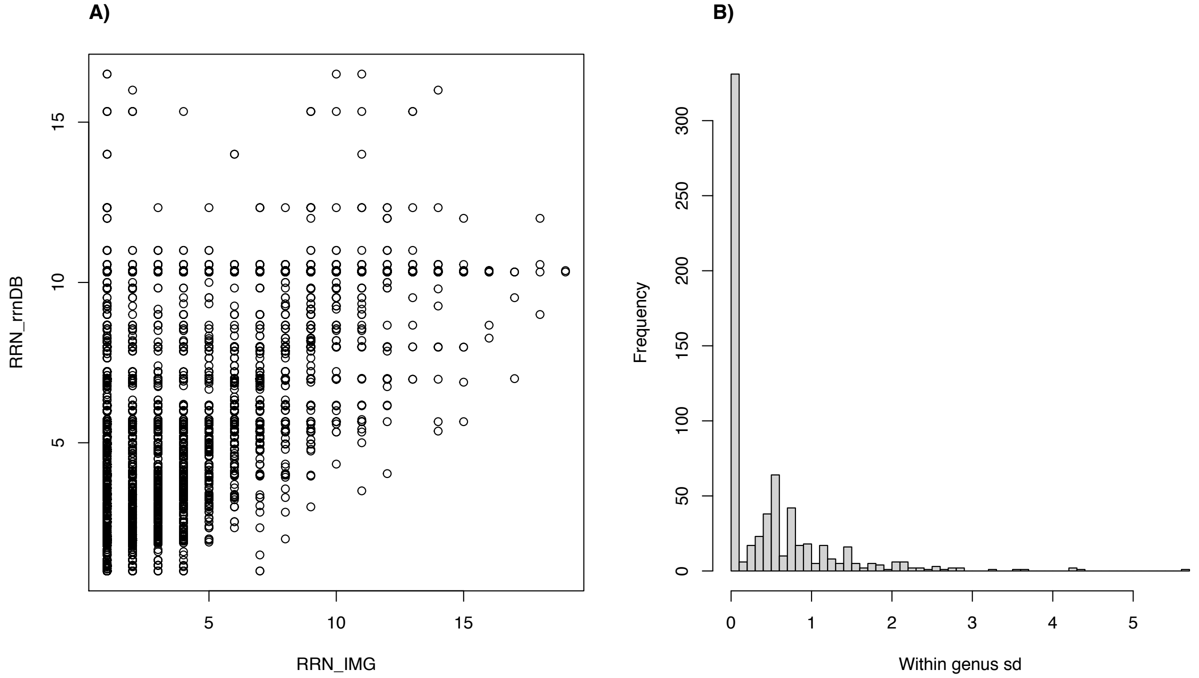


Figure S1: A) Correlation of RRN data from the JGI/IMG database (RRN_IMG) against RRN values that were extrapolated from genome level RRN values available via the rrnDB database indicate that a bias in the JGI/IMG values is mainly due to RRN underestimation for entries with high rrnDB RRN values. B) The standard deviation (sd) of within genus (NCBI taxonomy) RRN values for 666 genera of the rrnDB database with more than one entry per genus indicate values of sd < 1 in the large majority of genera.

Figure S2

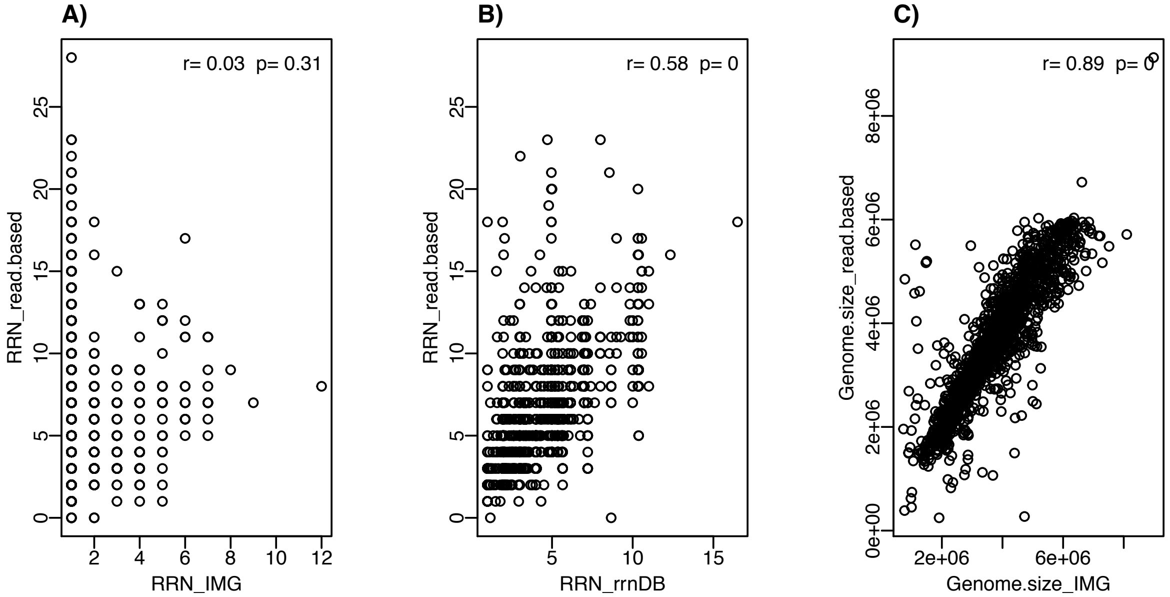


Figure S2: The JGI database provides information about NCBI accession, bioproject or biosample accession numbers, but no direct link to the raw read data. While for the majority of JGI entries listed in Table S1 it was not possible to identify runids by command-line based matches, we could extract raw read data for some of the entries. We applied the MicrobeCensus software to estimate the genome size as well as the number of sequenced genome equivalents based on the number of reads coding for single copy housekeeping genes relative to the number of total reads (Nayfach and Pollard, 2015). We excluded genomes with <2,000,000 sequenced reads and < 50 sequenced genome equivalents. We further applied the SortMeRNA software (Kopylova et al., 2012) to identify reads coding for the 16s rRNA and used the output in combination with the indicated number of sequenced genome equivalents to estimate the number of RRN per genome. A) Pearson correlation between the read based RRN estimate and the JGI RRN estimate. B) Pearson correlation between the read based RRN estimate and the rrnDB RRN estimate as given in Table S1. C) Pearson correlation between the read based genome size estimate and the genome size given in the JGI database.

While both the read-based and assembly-based estimation of RRN or genome size may suffer from biases, these biases are different and independent from each other. We therefore interpreted the strength of correlation between assembly-based and read-based estimates for RRN and genome size as measure for the quality of values provided in the rrnDB and JGI/IMG databases. The strongly improved correlation of read based RRN estimates against the rrnDB derived RRN values compared to a correlation against the JGI values, indicates that the presented rrnDB derived RRNs are exposed to reduced biases. However, the even better correlation between the genome size estimates in combination with low phylogenetic signals of RRN (Table 2, Figure 7) suggests that also the rrnDB derived RRN are likely still less accurate that are assembly-based genome estimates from the JGI/IMG database.

Figure S3

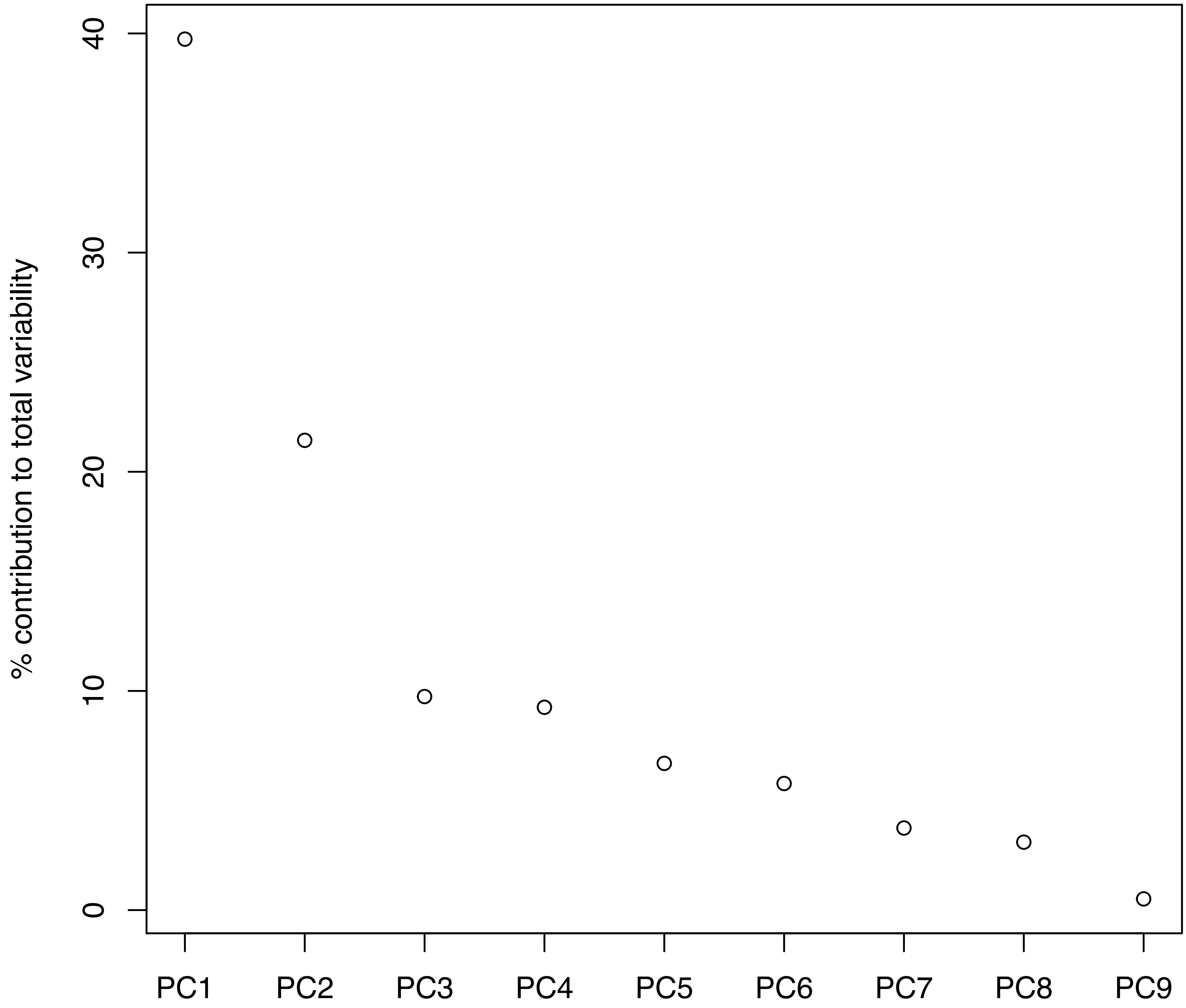


Figure S3: Percent contribution to variability of principal components PC1 to PC9.

### Figure S4

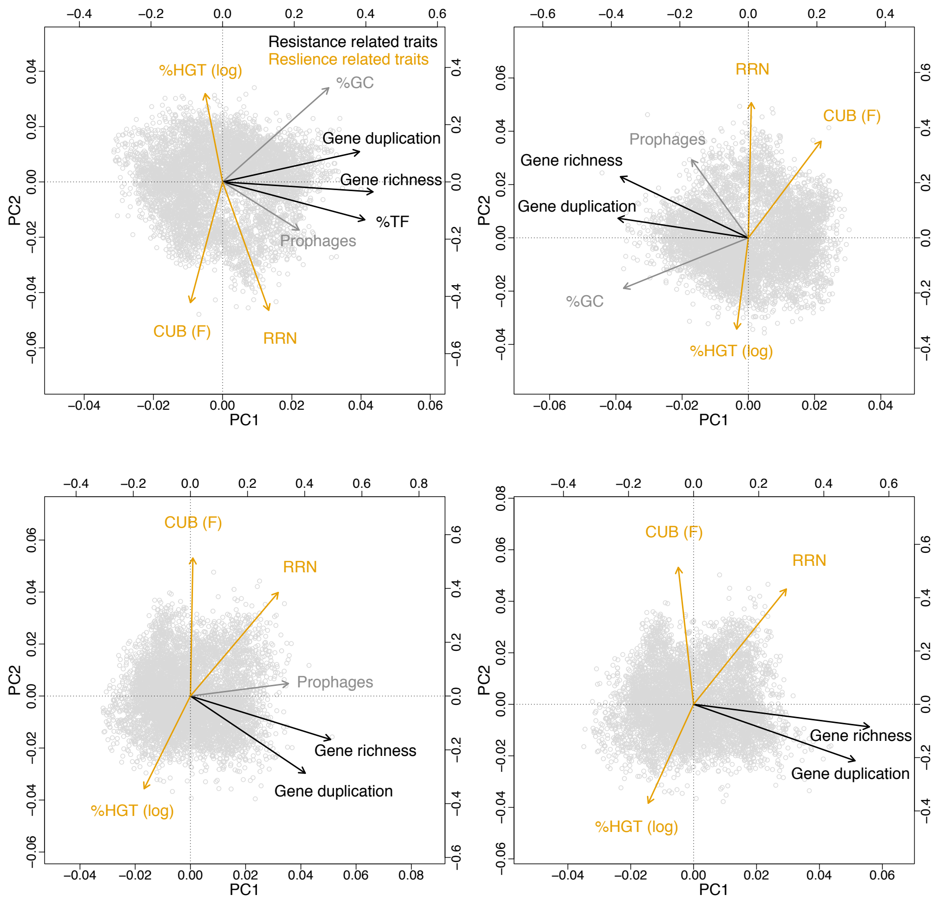


Figure S4: The random removal of 1,2,3 or 4 variable from the principal component analyses presented in Figure 3 demonstrates that the spatial patterns of resistance versus resilience related genomic traits are robust against the removal of individual variables from the dataset.

Table S1: Overview table for 17,856 genomes available via the JGI/IMG platform (https://img.jgi.doe.gov/, downloaded in August 2021), which are integrated in the reference database of the PICRUSt2 software (v2.1.2-b, Douglas et al., 2020). Some genomes present in the original PICRUSt2 reference database had meanwhile been replaced or removed. In the case of replaced genomes we indicate the new IMG genome ID in the column IMG.Genome.ID, while in the column PICRUSt.ID the former IMG genome ID is given. Douglas and colleagues furthermore present averaged information of several genomes, if these contained identical 16s rRNA genes (picrust.ID = *-cluster). However, the trait data presented in Table S1 refer exclusively to the genomes indicated in column IMG.Genome.ID.. All traits values were either directly extracted from the JGI/IMG database or computed with scripts that are available on GitHub (https://github.com/sarabeier/genomic.traits). RRN_IMG and RRN_rrnDB represent RRN values given by the JGI/IMG database and values extrapolated at the genus level (NCBI taxonomy) from the rrnDB, respectively. It was not possible to extract the information for all traits from all genomes and missing trait values are indicated by NA. Strains that could not be unambiguously binned at the species level via the GTDB taxonomy were classified via the FastANI software (Jain et al., 2018) into species level bins using an average nucleotide identity >94% (column FastANI_species). Information of all remaining columns is available via the JGI/IMG database.
